# A comparison of automatic cell identification methods for single-cell RNA-sequencing data

**DOI:** 10.1101/644435

**Authors:** Tamim Abdelaal, Lieke Michielsen, Davy Cats, Dylan Hoogduin, Hailiang Mei, Marcel J.T. Reinders, Ahmed Mahfouz

**Author notes:** Equal contribution (Tamim Abdelaal, Lieke Michielsen).

## Abstract

**Background:** Single cell transcriptomics are rapidly advancing our understanding of the cellular composition of complex tissues and organisms. A major limitation in most analysis pipelines is the reliance on manual annotations to determine cell identities, which are time-consuming and irreproducible. The exponential growth in the number of cells and samples has prompted the adaptation and development of supervised classification methods for automatic cell identification.

**Results:** Here, we benchmarked 20 classification methods that automatically assign cell identities including single cell-specific and general-purpose classifiers. The methods were evaluated using eight publicly available single cell RNA-sequencing datasets of different sizes, technologies, species, and complexity. The performance of the methods was evaluated based on their accuracy, percentage of unclassified cells, and computation time. We further evaluated their sensitivity to the input features, their performance across different annotation levels and datasets. We found that most classifiers performed well on a variety of datasets with decreased accuracy for complex datasets with overlapping classes or deep annotations. The general-purpose *SVM* classifier has overall the best performance across the different experiments.

**Conclusions:** We present a comprehensive evaluation of automatic cell identification methods for single cell RNA-sequencing data. All the code used for the evaluation is available on GitHub (https://github.com/tabdelaal/scRNAseq_Benchmark). Additionally, we provide a Snakemake workflow to facilitate the benchmarking and to support extension of new methods and new datasets (https://github.com/tabdelaal/scRNAseq_Benchmark/tree/snakemake_and_docker).

## Background

Single-cell transcriptomics (scRNA-seq) provides unprecedented opportunities to identify and characterize the cellular composition of complex tissues. Rapid and continuous technological advances over the past decade has allowed scRNA-seq technologies to scale to thousands of cells per experiment [1]. A common analysis step in analyzing single cell data involves the identification of cell populations presented in a given dataset. This task is typically solved by unsupervised clustering of cells into groups based on the similarity of their gene expression profiles, followed by cell population annotation by assigning labels to each cluster. This approach proved very valuable in identifying novel cell populations and resulted in cellular maps of entire cell lineages, organs and even whole organisms [2–7]. However, the annotation step is cumbersome and time-consuming as it involves manual inspection of cluster-specific marker genes. Additionally, manual annotations, which are often not based on standardized ontologies of cell labels, are not reproducible across different experiments within and across research groups. These caveats become even more pronounced as the number of cells and samples increases, preventing fast and reproducible annotations.

To overcome these challenges, a growing number of classification approaches are being adapted to automatically label cells in scRNA-seq experiments. scRNA-seq classification methods predict the identity of each cell by learning these identities from annotated training data (e.g. reference atlas). scRNA-seq classification methods are relatively new compared to the plethora of methods addressing different computational aspects of single cell analysis (e.g. normalization, clustering, and trajectory inference). However, the number of classification methods is rapidly growing to address the aforementioned challenges [8, 9]. While all scRNA-seq classification methods share a common goal, accurate annotation of cells, they differ in terms of their underlying algorithms and the incorporation of prior knowledge (e.g. cell type marker gene tables).

In contrast to the extensive evaluations of clustering, differential expression, and trajectory inference methods [10–12], there is currently only a single attempt comparing methods to assign cell type labels to cell clusters [13]. The lack of a comprehensive comparison of scRNA-seq classification methods leaves users without indications as to which classification method best fits their problem. More importantly, a proper assessment of existing approaches in comparison to baseline methods can greatly benefit new developments in the field and prevent unnecessary complexity.

Here, we benchmarked 20 classification methods to automatically assign cell identities including single cell-specific and general-purpose classifiers. The methods were evaluated using eight publicly available single cell RNA-sequencing datasets of different sizes, technologies, species, and complexity. The performance of the methods was evaluated based on their accuracy, percentage of unclassified cells, and computation time. We further evaluated their sensitivity to the input features, their performance across different annotation levels and datasets. In general, all classifiers perform well across all datasets, including the general-purpose classifiers. In our experiments, incorporating prior knowledge in the form of marker genes does not improve the performance. We observed large differences in the performance between methods in response to changing the input features. Furthermore, the tested methods vary considerably in their computation time which also vary differently across methods based on the number of cells and features. Our results highlight the general-purpose *SVM* classifier as the best performer overall.

## Results

### Benchmark of automatic cell identification methods

We benchmarked the performance and computation time of all 20 classifiers (Table 1) across all eight datasets (Table 2), whenever it is possible to apply. Classifiers can be divided into two categories: 1) supervised methods which require a training dataset labeled with the corresponding cell populations in order to train the classifier, or 2) prior-knowledge-supervised methods, for which either a marker genes file is required as an input, describing the signature genes to be expressed for each cell population, or a pre-trained classifier for specific cell populations is provided.

**Table 1.** Overview of the classification tools benchmarked during this study.

| Name | Version | Language | Type of Machine Learning | Prior knowledge | Rejection option | Reference |
| --- | --- | --- | --- | --- | --- | --- |
| Garnett | 0.1.4 | R | Generalized linear model | Yes | Yes | [23] |
| Moana | 0.1.1 | python | SVM with linear kernel | Yes | No | [24] |
| DigitalCellSorter | Github version: e369a34 | python | Voting based on cell type markers | Yes | No | [25] |
| SCINA | 1.1.0 | R | Bimodal distribution fitting for marker genes | Yes | No | [26] |
| scVI | 0.3.0 | python | Neural Network | No | No | [27] |
| Cell-Blast | 0.1.2 | python | Cell-to-cell similarity | No | Yes | [28] |
| ACTINN | GitHub version: 563bcc1 | python | Neural Network | No | No | [29] |
| Scmapcluster | 1.5.1 | R | Nearest median classifier | No | Yes | [17] |
| Scmapcell | 1.5.1 | R | kNN | No | Yes | [17] |
| scPred | 0.0.0.9000 | R | SVM with radial kernel | No | Yes | [30] |
| CHETAH | 0.99.5 | R | Correlation to training set | No | Yes | [31] |
| CaSTLe | Github version: 258b278 | R | Random Forest | No | No | [32] |
| SingleR | 0.2.2 | R | Correlation to training set | No | No | [33] |
| scID | 0.0.0.9000 | R | LDA | No | Yes | [34] |
| singleCellNet | 0.1.0 | R | Random Forest | No | No | [35] |
| LDA | 0.19.2 | python | LDA | No | No |  |
| NMC | 0.19.2 | python | NMC | No | No |  |
| RF | 0.19.2 | python | RF (50 trees) | No | No |  |
| SVM | 0.19.2 | python | SVM (linear kernel) | No | No |  |
| kNN | 0.19.2 | python | kNN (k = 50) | No | No |  |

**Table 2.**
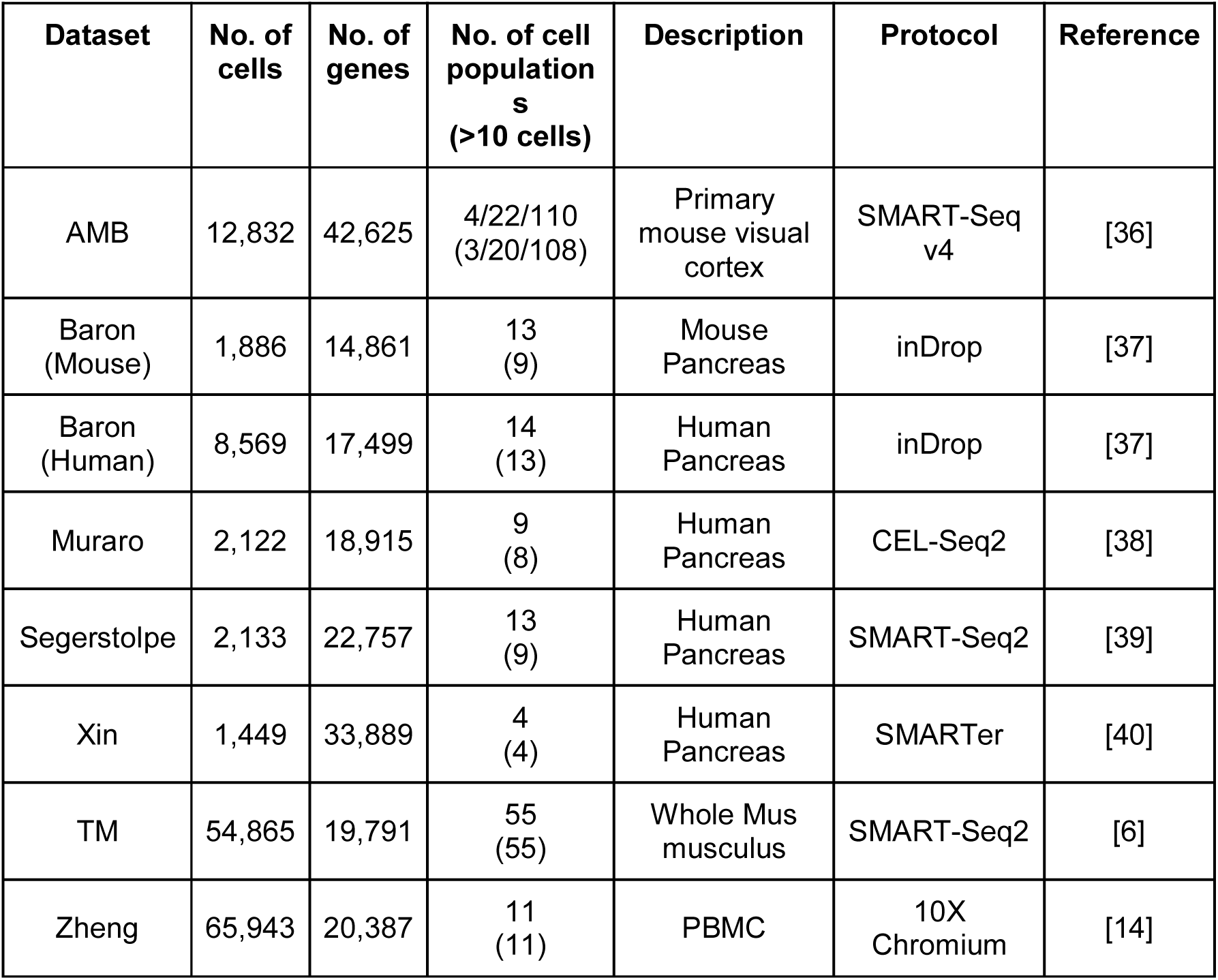
Overview of the datasets used during this study.

The datasets used in this study vary in the number of cells, genes and cell populations (annotation level), in order to represent different levels of challenges in the classification task and to evaluate how each classifier performs in each case (Table 2). Starting from relatively typical sized scRNA-seq datasets (∼1,500 – ∼8,500 cells), such as the five pancreatic datasets (Baron Mouse and Human, Muraro, Segerstolpe and Xin), which include both mouse and human pancreatic cells and vary in the sequencing protocol used. The Allen Mouse Brain (AMB) dataset is used to evaluate how the classification performance changes when dealing with different levels of cell population annotation since the AMB dataset contains three levels of annotations for each cell (3, 20 or 108 cell populations), denoted as AMB3, AMB20, and AMB108. The Tabula Muris (TM) and Zheng datasets represent relatively large scRNA-seq datasets (>50,000 cells), to assess how well the classifiers scale with large datasets. Additionally, by including the Zheng dataset, we are able to benchmark four prior-knowledge-supervised classifiers, since the marker genes files or pre-trained classifier are available for the four classifiers for peripheral blood mononuclear cells (PBMCs).

Due to either CPU time constraint or memory requirement of some classifiers, it was not possible to apply them on the large datasets, e.g., TM and Zheng. *Cell-BLAST* requires a lot of memory (> 100 GB) and long run time (in order of days) to obtain predictions for ∼10,000 cells, and *SingleR* has long computation time similar to *Cell-BLAST*. Therefore, we did not evaluate *Cell-BLAST* on the TM and Zheng datasets, and *SingleR* was not evaluated on the Zheng dataset. Moreover, *scPred* failed while being tested on the Zheng dataset.

### Overall performance evaluation across datasets and methods

Generally, all classifiers perform well across all datasets, including the general-purpose classifiers (Figure 1), except *Cell-BLAST* which had remarkably lower performance compared to all other classifiers across all datasets. Further, *scVI* has low performance on the deeply annotated datasets TM (55 cell populations) and AMB108 (108 cell populations), and *kNN* produces low performance for the Xin and AMB108 datasets.

**Figure 1.**
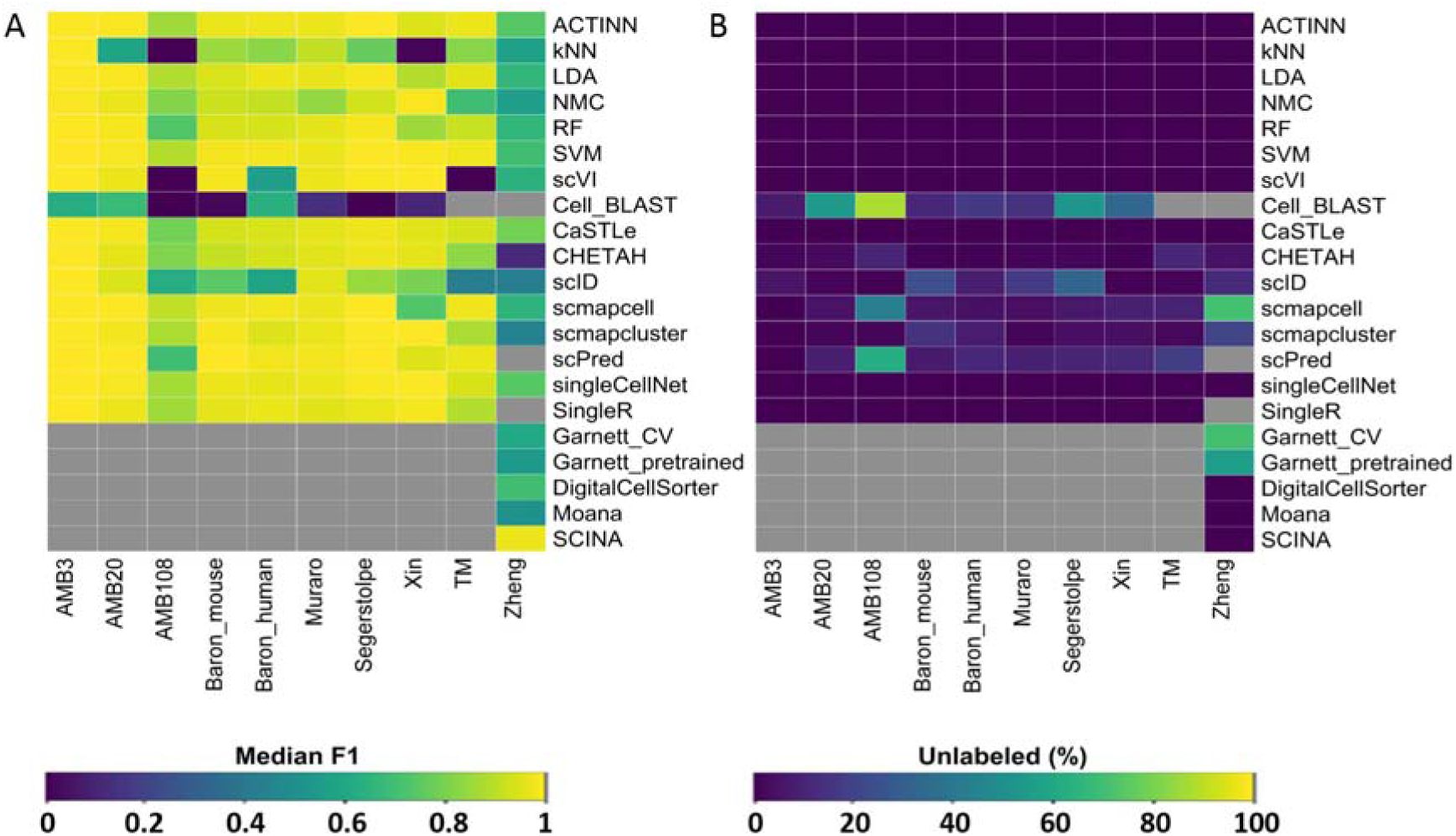
Performance comparison of supervised classifiers for cell identification using different scRNA-seq datasets. **(A)** Heatmap of the median F1-score across all cell populations per classifier per dataset. **(B)** Percentage of unlabeled cells across all cell populations per classifier per dataset. Light-grey boxes indicate that the corresponding method could not be tested on the corresponding dataset.

For the pancreatic datasets, the best-performing classifiers are *SVM, scPred, scmapcell, scmapcluster, scVI, ACTINN, singleCellNet, LDA* and *NMC. SVM* is the only classifier to be in the top five list for all five pancreatic datasets, while *NMC*, for example, appears only in the top five list for the Xin dataset. The Xin dataset contains only four major pancreatic cell types (alpha, beta, delta and gamma) making the classification task relatively easy for all classifiers, including *NMC*. Considering only the median F1-score can be misleading since some classifiers incorporate a rejection option (e.g. *scmapcell, scPred*), by which a cell is assigned as ‘unlabeled’ if the classifier is not confident enough. Figure 1B summarizes the percentage of unlabeled cells for each classifier. In the Baron Human dataset, for example, the median F1-score for *scmapcell, scPred* and *SVM* is 0.984, 0.981, and 0.980, respectively. However, *scmapcell* and *scPred* assigned 4.2% and 10.8% of the cells, respectively, as unlabeled while *SVM* classified 100% of the cells. This shows an overall better performance for *SVM*.

For the TM dataset, the top five performing classifiers are *SVM, scmapcell, scPred, ACTINN* and *LDA* with a median F1-score > 0.95, showing that these classifiers can perform well and scale to large scRNA-seq datasets with a deep level of annotation. Furthermore, *scmapcell* and *scPred* assigned 9.5% and 17.7% of the cells as unlabeled, which shows a superior performance for *SVM* with high F1-score and no unlabeled cells.

### Incorporating marker-genes does not improve performance on PBMC data

For the Zheng dataset, *Garnett, Moana, DigitalCellSorter* and *SCINA* could be evaluated and benchmarked with the rest of the classifiers. Although the best performing classifier is *SCINA* with a median F1-score of 0.968, this performance is based only on 3, out of 11, cell populations (Monocytes, B cells and NK cells) for which marker-genes are provided. Supplementary Table 1 summarizes which cell populations from the Zheng dataset can be classified by the prior-knowledge-supervised methods. Interestingly, none of the prior-knowledge-supervised methods showed superior performance compared to other classifiers. Beside *SCINA*, the top classifiers are *CaSTLe, ACTINN, singleCellNet* and *SVM*. Generally, all classifiers show relatively lower performance on the Zheng dataset compared to other datasets, as the Zheng dataset contains 11 immune cell populations which are harder to differentiate, particularly the T cell compartment (6 out of 11 cell populations). This difficulty of separating these populations was previously noted in the original study [14]. Also, the confusion matrices for *CaSTLe, ACTINN, singleCellNet* and *SVM* clearly indicate that some populations are similar to each other where all classifiers are making wrong predictions, such as 1) monocytes with dendritic cells, 2) two CD8+ T populations, and 3) four CD4+ T populations (Supplementary Figure 1).

### Performance evaluation across different annotation levels

We used the AMB dataset with its three different levels of annotations, to evaluate the classifiers’ performance behavior with a larger number of smaller cell populations within the same dataset. For AMB3, the classification task is relatively easy, differentiating between three major brain cell types (GABAergic, Glutamatergic and Non-Neuronal). All classifiers perform almost perfectly with a median F1-score > 0.99, except *Cell-BLAST* (median F1-score = 0.619) (Figure 1). For AMB20, the classification task becomes slightly more challenging and the performance of some classifiers drops, especially *kNN*. The low performance of *kNN* in this case is due to the setting of *k* = 50, a parameter we did not optimize, which is larger than the size of three out of 20 classes leading to misclassifications by *kNN*. The top five classifiers are *scmapcell, scPred, SVM, ACTINN* and *LDA*, where *scmapcell* and *scPred* assigned 4.9% and 8.4% of the cells as unlabeled. For the deeply annotated AMB108 dataset, the performance of all classifiers drops further, except for *kNN* and *scVI*, where the median F1-score is zero. The top five classifiers are *scmapcell, SVM, LDA, scmapcluster* and *singleCellNet*, with *scmapcell* assigning 41.9% of the cell as unlabeled. These results show an overall superior performance for general-purpose classifiers (*SVM* and *LDA*) compared to other scRNA-seq specific classifiers across different levels of cell population annotation.

Instead of only looking at the median F1-score, we also evaluated the F1-score per cell population for each classifier (Figure 2). We confirmed previous conclusions, *Cell-BLAST* exhibits low performance in general (Figure 2A-C), *kNN* performance drops with deep annotation having smaller cell populations (Figure 2B-C), *scVI* poorly performs on the deeply annotated AMB108 dataset. Additionally, we could observe that some cell populations are much harder to classify compared to other populations, for example Serpinf1 cells in the AMB20 dataset.

**Figure 2.**
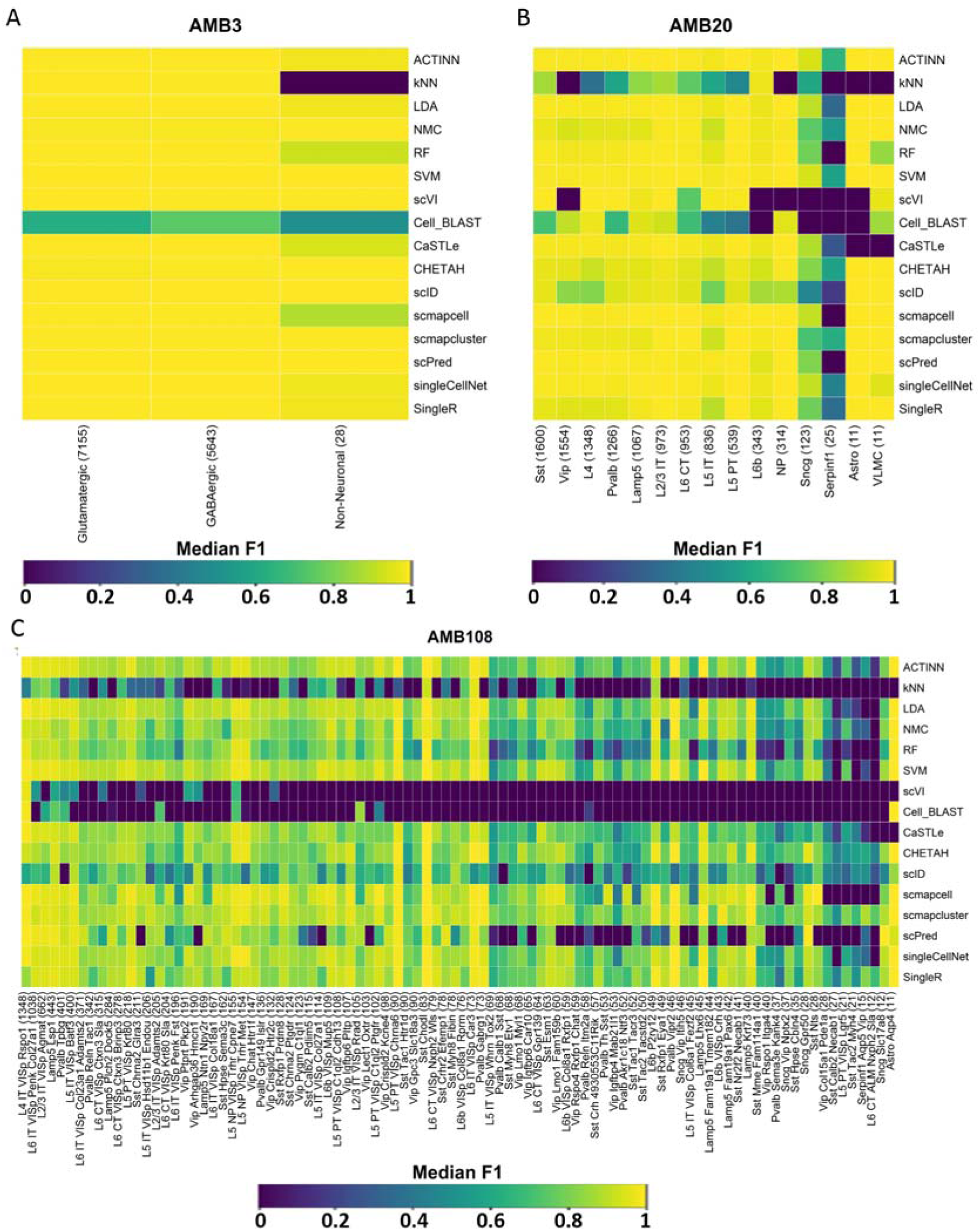
Classification performance across different annotation levels in the Allen Mouse Brain dataset. Heatmaps show the F1-scores of each method for each cell population in the **(A)** AMB3, **(B)** AMB20, and **(C)** AMB108 datasets. The cell populations are sorted from left-to-right in descending order according to their size (i.e. number of cells). The size of each population is indicated between brackets.

### Performance evaluation across datasets

While evaluating the classification performance within a dataset is important, it is more challenging to predict cell identities across datasets. To test the classifiers’ ability to predict cell identities in a dataset that was not used for training, we used the four human pancreatic datasets: Baron Human, Muraro, Segerstople and Xin. In this case, the classification performance can be affected by batch differences between datasets. We evaluated the performance of the classifiers when trained using the raw data as well as aligned data using the mutual nearest neighbor (MNN) method [15]. Supplementary Figure 2 shows UMAPs [16] of the combined dataset before and after alignment, demonstrating better grouping of pancreatic cell types after alignment.

For the raw (unaligned) data, the best performing classifiers across all four datasets are *SVM, scVI, scmapcell, ACTINN* and *singleCellNet* (Figure 3A,C). For the aligned data, the best performing classifiers are *SVM, singleCellNet, kNN* and *NMC* (Figure 3B,D). Some classifiers benefit from aligning the datasets such as *kNN, NMC* and *singleCellNet*, producing higher median F1-scores (Figure 3A,B). On the other hand, some other classifiers failed the classification task completely, such as *scmapcell* which labels all cells as unlabeled. Some other classifiers failed to run over the aligned datasets, such as *ACTINN, scVI, Cell-BLAST, scID, scmapcluster* and *scPred*. These classifiers work only with positive gene expression data, while the aligned datasets contains positive and negative gene expression values.

**Figure 3.**
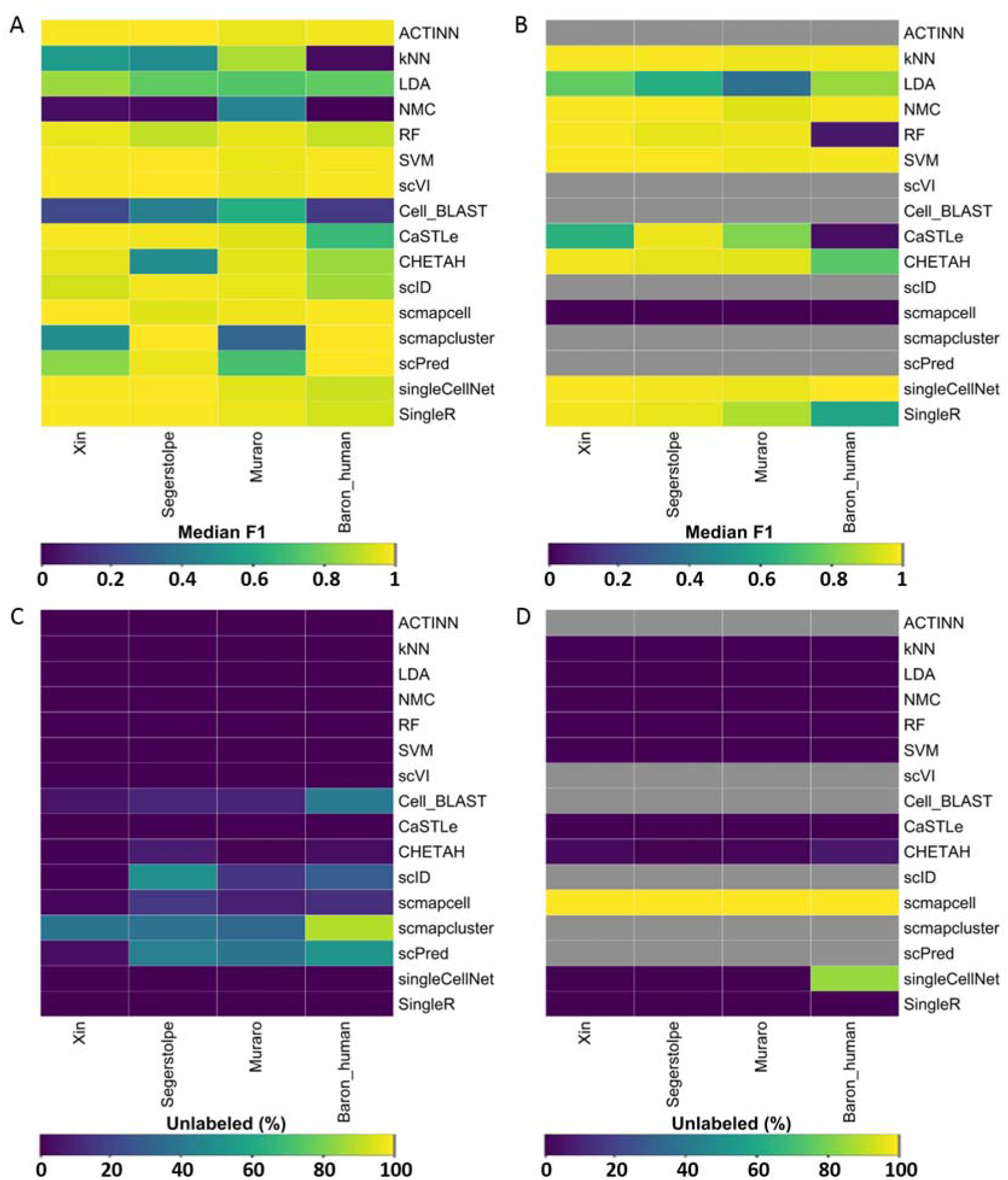
Classification performance across different pancreatic datasets. Heatmaps showing **(A-B)** the median F1-score and **(C-D)** the percentage of unlabeled cells for each classifier. **(A,C)** Show the results for the unaligned datasets. **(B,D)** Show the results for the aligned datasets using MNN. The column labels indicates which of the four datasets was used as a test set, in which case the other three sets were used to train the classifiers. Light-grey boxes indicate that the corresponding method could not be tested on the corresponding dataset.

### Performance sensitivity to the input features

During the cross-validation experiment described earlier, we used all features (genes) as input to the classifiers. However, some classifiers suffer from overtraining when too many features are used. Therefore, we tested the effect of feature selection on the performance of the classifiers. Different strategies for feature selection in scRNA-seq classification experiments exist. Using genes as features that have a higher number of dropouts compared to the expected number of dropouts has been shown to yield the best results [17, 18]. Here, subsets of features were selected based on this criterion. The feature selection experiments were all done on the TM dataset. For the number of features, we used the top: 100, 200, 500, 1000, 2000, 5000, and 19791 (all) genes. Some classifiers include a built-in feature selection method which is used by default. To ensure that all tools use the same set of features, the built-in feature selection was turned off during these experiments. Due to long running times or excessive memory usage, not all feature sets could be tested for all tools. As already discussed before, *Cell-BLAST* could not be tested on the TM dataset. During feature selection, we ran *Cell-Blast* on all feature sets except the largest set with all features. *scVI* also timed out when running on this feature set. Furthermore, *scPred* failed when tested using 2000 features, and *singleCellNet* timed out when tested using 5000 features.

Figure 4 presents the performance of the classifiers using the different sets of features. Some methods are clearly overtrained when the number of features increases. *scmapcell*, for instance, shows the highest median-F1 score when using less features, but its performance drops when the number of features increases. On the contrary, the performance of some classifiers, such as *SVM*, keeps improving when the number of features increases. These results indicate that the optimal number of features is different for each classifier.

**Figure 4.**
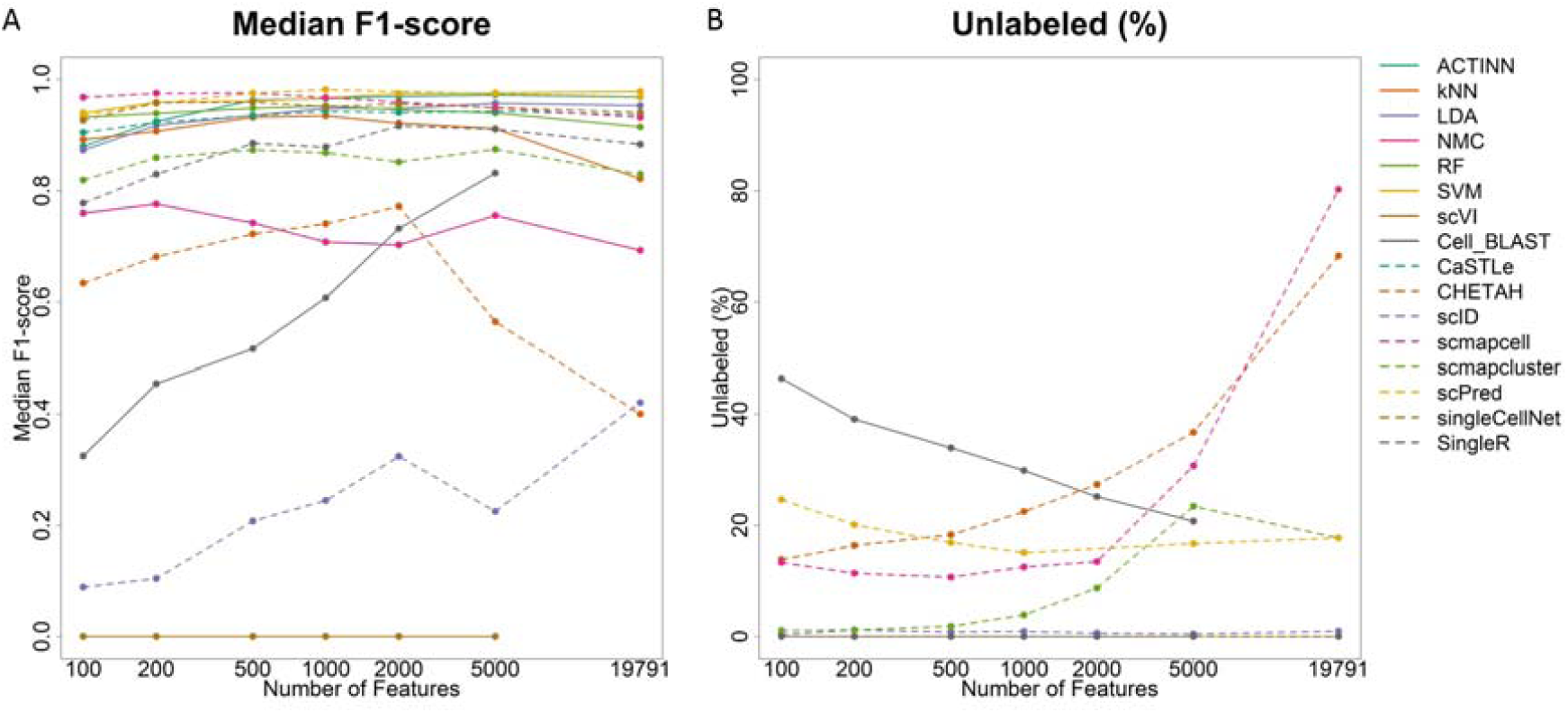
Classification performance across different number of features. Line plots show **(A)** the median F1-score and (**B**) percentage of unlabeled cells of each classifier applied to the TM dataset with the top 100, 200, 500, 1000, 2000, 5000, and 19791 (all) genes as input feature sets. Genes were ranked based on dropout-based feature selection. The x-axis is log-scaled in all panels. Few points are not shown as the corresponding classifier failed or timed out when tested.

Looking at the median-F1 score, there are several methods with a high performance. *ACTINN, LDA, RF, scmapcell, scPred, singleCellNet*, and *SVM* all have a median F1-score higher than 0.95 for one or more of the feature sets. Some of these well-performing tools, however, leave many cells unlabeled. *scmapcell*, for instance, yields a median F1-score of 0.976 when using a subset of 500 genes, with 10% of the cells is still unlabeled. The same holds for *scPred*, overall, it has the highest median F1-score (0.982) when using 1000 genes, with 15% of the cells remains unassigned. *ACTINN, SVM, LDA, singleCellNet*, and *RF*, on the contrary label all the cells. Overall *SVM* shows the second highest performance with a score of 0.979. It thus performs slightly worse than *scPred*, but it does label all the cells.

### Running time evaluation

To compare the runtimes of the tools and see how they scale when the number of cells increases, we compared the number of cells in each dataset with the computation time of the tools (Figure 5A). Overall, big differences in the computation time can be observed when comparing the different methods. For example, for the Zheng dataset the runtime varies between 9.65 seconds for *scmapcluster* and 6.00 hours for *LDA. singeCellNet* showed the longest computation time overall. Running *singleCellNet* on the TM dataset took more than 25 hours. In general, all tools show an increase in computation time when the number of cells increase. However, when comparing the largest datasets, TM and Zheng, not all tools show an increase in computation time. Despite the increase in the number of cells between the datasets, *CaSTLe, CHETAH*, and *SingleR*, have a decreasing computation time. A possible explanation could be that the runtime of these tools also depends on the number of genes or the number of cell populations in the dataset. The Zheng dataset, for example, contains less cell populations than the TM dataset (11 compared to 55). To evaluate this properly, the runtime of the tools on the AMB3, AMB20, and AMB108 datasets were compared (Figure 5B), and this shows an increase in run time when the number of cell populations increases, while the number of cells and genes remains constant. For other tools, such as *ACTINN* and *scmapcell*, the runtime does not increase.

**Figure 5.**
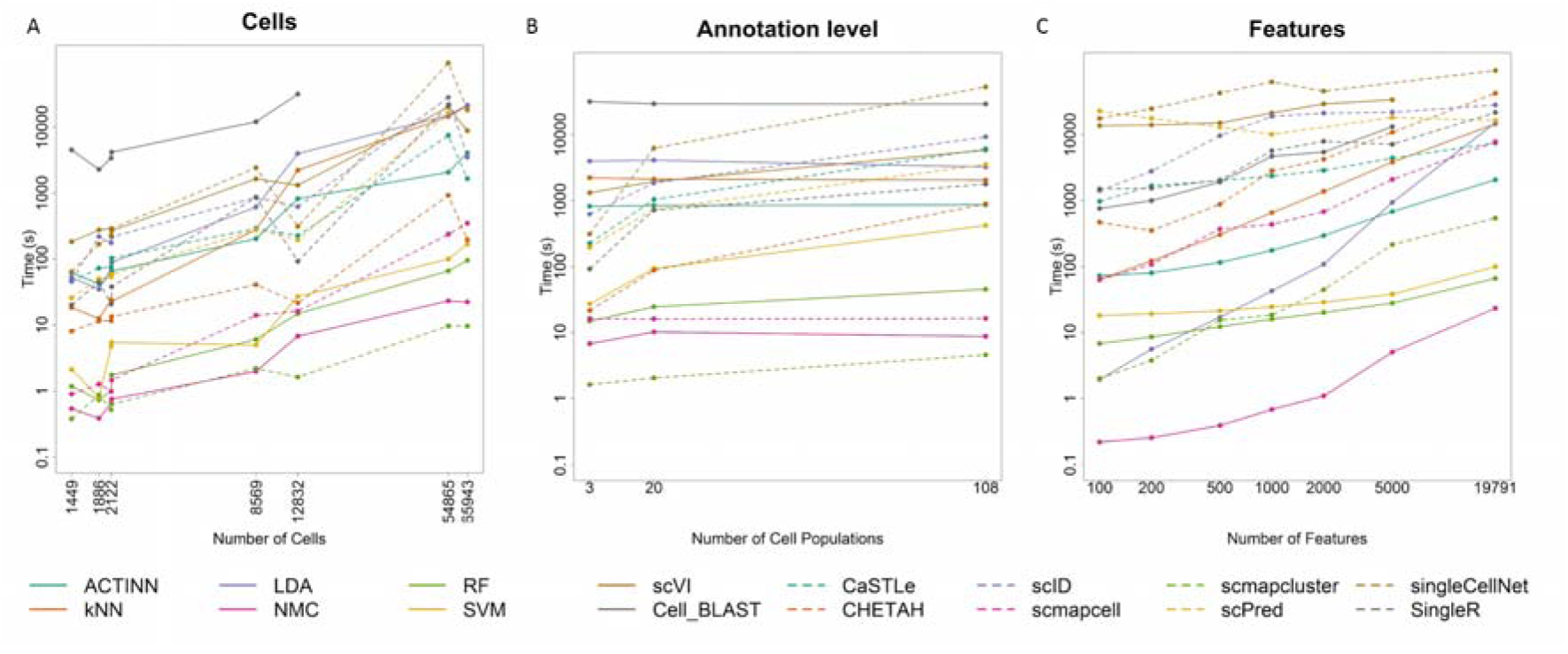
Computation time Evaluation. The computation time of each tool is plotted against **(A)** the number of cells, **(B)** the number of cell populations, and **(C)** the number of features (genes). The x-axis is log-scaled in all panels. Few points are not shown as the corresponding classifier failed or timed out when tested.

Some of the tools even have a high runtime for the small datasets. On the smallest, Xin, dataset all classifiers have a computation time < 5 minutes, with most classifiers finishing within 60 seconds. *Cell-BLAST*, however, takes more than 75 minutes.

To assess the effect of the number of genes on the computation time, we compared the computation time of the methods during the feature selection experiments (Figure 5C). Most methods scale linearly with the number of genes. However, *LDA* does not scale very well when the number of genes increases. If the number of features is higher than the number of cells, the complexity of *LDA* is O(*g*^3), where *g* is the number of genes [19]. The computation time of the methods is thus dependent on the number of cells, number of genes, and, for most tools, also the number of different cell populations in the dataset.

Five classifiers, *scmapcell, scmapcluster, SVM, RF*, and *NMC*, have a computation time below six minutes on all the datasets. Here, it is especially noteworthy that most of these tools, and *SVM* in particular, also have the highest median F1-scores during all previous experiments.

## Discussion

In this study, we evaluated the performance of 20 different methods for automatic cell identification using eight scRNA-seq datasets. Several classifiers accurately performed on almost all datasets, particularly: *SVM, scPred, scmapcell/cluster, singleCellNet, scVI, LDA* and *ACTINN*. Considering all three evaluation metrics (median F1-score, % of unlabeled cells and computation time), *SVM* is overall the best performing classifier for the scRNA-seq datasets used. Our results show that *SVM* scales well to large datasets as well as deep annotation levels. In addition, *SVM* did not suffer from the large number of features (genes) present in the data, producing the highest performance on the TM dataset using all genes, due to the incorporated L2-regularization. The comparable or higher overall performance of a general-purpose classier such as *SVM* warrants caution when designing scRNA-seq specific classifiers that they do not introduce unnecessary complexity.

*scPred*, which is based on a *SVM* with radial kernel, performed well on most dataset, yet it suffers from long computation time for large datasets, together with *LDA, ACTINN* and *singleCellNet*, where the latter becomes even slower with large number of cell populations. In addition, in some cases, *scPred* and *scmapcell/cluster* reject high proportions of cells as unlabeled. In general, incorporating a rejection option with classification is a good practice, as it allows to detect potentially new cell populations not included in the training data, and improve the performance for the classified cells with high confidence. However, for the datasets used in this study, the performance of classifiers with rejection option did not show substantial improvement compared to other classifiers. *scVI* works well for datasets with relatively small number of cell populations, but failed to scale with deeply annotated datasets. *kNN* classifier produces poor performance with most datasets, but this performance can potentially be improved by optimizing the number of neighbors. Generally, we evaluated all classifiers using their default settings. However, adjusting these settings for a specific dataset might improve the performances but increases the risk of overtraining.

For the Zheng dataset, the prior-knowledge-supervised methods did not improve the classification performance over supervised methods which do not incorporate such prior knowledge. These results indicate that incorporating prior knowledge in the form of marker genes is not beneficial. Besides, defining these marker genes is often challenging and heavily depends on personal expertise. Furthermore, these marker genes can be implicitly learned by supervised methods through the training process.

Based on our results, we recommend to use of the general-purpose *SVM* classifier (with a linear kernel) since it had better or equal performance compared to the other classifiers tested across all datasets, with a remarkably fast computation time. Other high performing classifiers include: *scPred, scmapcell/cluster, singleCellNet, LDA* and *ACTINN*. While the performance of almost all methods was relatively high on various datasets, some datasets with overlapping populations (e.g. Zhang PBMC dataset) remain challenging.

## Conclusions

We present a comprehensive evaluation of automatic cell identification methods for single cell RNA-sequencing data. Generally, all classifiers perform well across all datasets, including the general-purpose classifiers. In our experiments, incorporating prior knowledge in the form of marker genes does not improve the performance (on PBMC data). We observed large differences in the performance between methods in response to changing the input features. Furthermore, the tested methods vary considerably in their computation time which also vary differently across methods based on the number of cells and features. Our results highlight the general-purpose *SVM* classifier as the best performer overall. To support future extension of this benchmarking work with new classifiers and datasets, we provide a Snakemake workflow to automate the performed benchmarking analyses (https://github.com/tabdelaal/scRNAseq_Benchmark/tree/snakemake_and_docker).

## Methods

### Classification methods

We evaluated 20 scRNA-seq classifiers, publicly available as R or Python packages or scripts (Table 1). This set included 15 methods developed specifically for scRNA-seq data as well as 5 general-purpose classifiers from the scikit-learn library in Python: linear discriminant analysis (*LDA*), nearest mean classifier (*NMC*), k-nearest neighbor (*kNN*), support vector machine (*SVM*), and random forest (*RF*). Methods were excluded from the evaluation if they did not return the predicted labels for each cell. For example, we excluded *LAmbDA [20]* because the tool only returns the posterior probabilities rather than predicted labels. Similarly, we excluded *MetaNeighbor [21]* because the tool only returns the area under the receiver operator characteristic curve (AUROC). For all tools the latest (May 2019) package was installed or scripts were downloaded from their Github. For *scPred* it should be noted that it is only compatible with an older version of Seurat (v2.0). For *CHETAH* it is important that the R version 3.6 or newer is installed.

During the benchmark, all tools were run using their default settings and if not available, we used the settings provided in the accompanying examples and vignettes. As input, we provided each method with the raw count data (after cell and gene filtering as described in Data Preprocessing) according to the method documentation. The majority of the methods have a built-in normalization step. For the general-purpose classifiers, we provided log-transformed counts, *log*_2_ (*count* + 1).

Some methods required a marker gene file as an input (e.g. *Garnett, Moana, SCINA, DigitalCellSorter*). In this case, we use the marker gene files provided by the authors. We did not attempt to include additional marker gene files and hence the evaluation of those methods is restricted to datasets where a marker gene file for cell populations is available.

### Datasets

Eight scRNA-seq datasets were used to evaluate and benchmark all classification tools (Table 2). Datasets vary across species (human and mouse), tissue (brain, pancreas, PBMC and whole mouse), as well as the sequencing protocol used. The Allen Mouse Brain (AMB) dataset was downloaded from http://celltypes.brain-map.org/rnaseq. All five pancreatic datasets were obtained from: https://hemberg-lab.github.io/scRNA.seq.datasets/. The Tabula Muris (TM) dataset was downloaded from https://tabula-muris.ds.czbiohub.org/. For the PBMC dataset, we downloaded the gene-cell count matrix for the ‘Fresh 68k PBMCs’ [14] from: https://support.10xgenomics.com/single-cell-gene-expression/datasets. The cell population annotation for all datasets was provided with the data, except the Zheng dataset, for which we obtained the cell population annotation from https://github.com/10XGenomics/single-cell-3prime-paper/tree/master/pbmc68k_analysis. These annotations were used as ‘ground truth’ during the evaluation of the cell population prediction obtained from the classification tools.

### Data Preprocessing

Based on the manual annotation provided in the datasets, we started by filtering out cells that were labeled as doublets, debris or unlabeled cells. Next, we filtered genes with zero counts across all cells. For cells, we calculated the median number of detected genes per cell, and from that we obtained the median absolute deviations (MADs) across all cells in the log scale. We filtered out cells when the total number of detected genes was below 3 MADs from the median number of detected genes per cell. The number of cells and genes in Table 2 represent the size of each dataset after this stage of preprocessing.

Moreover, before applying cross validation to evaluate each classifier, we excluded cell populations with less than 10 cells across the entire dataset, Table 2 summarizes the number of cell populations before and after this filtration step for each dataset.

### Experimental setup

For the supervised classifiers, we evaluated the performance by applying a 5-fold cross validation across each dataset after filtering genes, cells and small cell populations. The folds were divided in a stratified manner in order to keep equal proportion of each cell population in each fold. The training and test indices for each fold were defined and saved for each dataset, these indices were provided while applying the classifiers on the datasets, to make sure all folds are exactly the same for all classifiers.

The prior-knowledge-supervised classifiers, *Garnett, Moana, DigitalCellSorter* and *SCINA*, were only evaluated on the Zheng dataset, for which the marker genes file or the pre-trained classifier was available, after filtering genes and cells. Each classifier uses the dataset and the marker genes file as inputs, and outputs the cell population label corresponding to each cell. No cross validation is applied in this case, except for *Garnett* where we could either use the pre-trained version provided from the original study, or train our own classifier using the marker genes file along with the training data. In this case, we applied 5-fold cross validation using the same train and test indices described previously. Supplementary Table 1 shows the mapping of cell populations between the Zhang dataset and each of the prior-knowledge-supervised classifiers. For *Moana* a pre-trained classifier was used, this classifier also predicted cells to be Memory CD8+ T cells and CD16+ Monocytes, while these cell populations were not in the Zheng dataset.

### Across dataset prediction

We selected the major four endocrine pancreatic cell types (alpha, beta, delta and gamma) across all four human pancreatic datasets: Baron Human, Muraro, Segerstolpe and Xin. Supplementary table 2 summarizes the number of cells in each cell type across all datasets. To account for batch effects and technical variations between different protocols, datasets were aligned using MNN [15] from the scran R package (version 1.1.2.0). Using both the raw data (unaligned) and the aligned data, we applied leave-one-dataset-out cross validation where we train on three datasets and test on the left out dataset.

### Performance evaluation metrics

The performance of the tools on the datasets is evaluated using three different metrics: 1) For each cell population in the dataset the F1-score is reported. The median of these F1-scores is used as a measure for the performance on the dataset. 2) Some of the tools do not label all the cells. These unassigned cells are not considered in the F1-score calculation. The percentage of unlabeled cells is also used to evaluate the performance. 3) The computation time of the tools is also measured.

### Feature selection

Genes are selected as features based on their dropout rate. The method used here, is based on the method described in [17]. During feature selection, a sorted list of the genes is made. Based on this list, the top *n* number of genes can be easily selected during the experiments. First, the data is normalized using *log*_2_ (*count* + 1). Next, for each gene the percentage of dropouts, *d*, and the mean, *m*, of the normalized data are calculated. Genes that have a mean or dropout rate of zero are not considered during the next steps. These genes will be at the bottom of the sorted list. For all other genes, a linear model is fitted to the mean and log2(*d*). Based on their residuals, the genes are sorted in descending order and added to the top of the list.

### Benchmarking pipeline

In order to ensure reproducibility and support future extension of this benchmarking work with new classification methods and benchmarking datasets, a Snakemake [22] workflow for automating the performed benchmarking analyses was developed with an MIT license (https://github.com/tabdelaal/scRNAseq_Benchmark/tree/snakemake_and_docker). Each tool (license permitting) is packaged in a Docker container (https://hub.docker.com/u/scrnaseqbenchmark) alongside the wrapper scripts and their dependencies. These images will be used through snakemake’s singularity integration to allow the workflow to be run without the requirement to install specific tools and to ensure reproducibility. Documentation is also provided to execute and extend this benchmarking workflow to help researchers to further evaluate interested methods.

## Declarations

### Ethics approval and consent to participate

Not applicable.

### Consent for publication

Not applicable.

### Availability of data and material

The filtered datasets analyzed during the current study are available in the Zenodo repository [https://zenodo.org/record/2877646#.XN8l_kzapo] Code: https://github.com/tabdelaal/scRNAseq_Benchmark

## Authors’ contributions

TA, LM, MJTR, and AM conceived the study and designed the experiments. TA and LM performed the experiments. DH, DC, and HM designed and developed the Snakemake workflow. MJTR and AM supervised the experiments. TA, LM, HM, and AM wrote the manuscript. All authors reviewed and approved the manuscript.

## Acknowledgements

Not applicable.

## Supplementary Materials

**Supplementary Table 1.** Mapping of true cell population labels from the Zheng dataset to cell population labels of the prior-knowledge-supervised classifiers.

| True Labels | Garnett | DigitalCellSorter | Moana | SCINA |
| --- | --- | --- | --- | --- |
| CD14+ Monocyte | CD14+ Monocyte | CD14+ Monocyte | CD14+ Monocyte | CD14+ Monocyte |
| Dendritic | Dendritic | Dendritic | Dendritic |  |
| CD34+ | CD34+ |  |  |  |
| CD56+ NK | CD56+ NK | CD56+ NK | CD56+ NK | CD56+ NK |
| CD19+ B | CD19+ B | CD19+ B | CD19+ B | CD19+ B |
| CD4+ T Helper 2 | CD4+ T cell | T cell |  |  |
| CD4+/CD25 T Reg |  |  |  |  |
| CD4+/CD45RA+/CD25- Naïve T |  |  | Naïve CD4+ T cells |  |
| CD4+/CD45RO+ Memory |  |  | Memory CD4+ T cells |  |
| CD8+ Cytotoxic T | CD8+ T cell | T cell |  |  |
| CD8+/CD45RA+ Naïve Cytotoxic |  |  | Naïve CD8+ T cells |  |
|  |  |  | Memory CD8+ T cells |  |
|  |  |  | CD16+ Monocytes |  |

**Supplementary Table 2.** Cell type size for each pancreatic dataset used in the across dataset performance evaluation.

| **Dataset** | **alpha** | **beta** | **delta** | **gamma** | **Total** |
| --- | --- | --- | --- | --- | --- |
| **Baron (Human)** | 2326 | 2525 | 601 | 255 | **5707** |
| **Muraro** | 812 | 448 | 193 | 101 | **1554** |
| **Segerstolpe** | 872 | 263 | 110 | 195 | **1440** |
| **Xin** | 855 | 466 | 46 | 82 | **1449** |
| **Total** | **4865** | **3702** | **950** | **633** | **10150** |

**Supplementary Figure 1.**
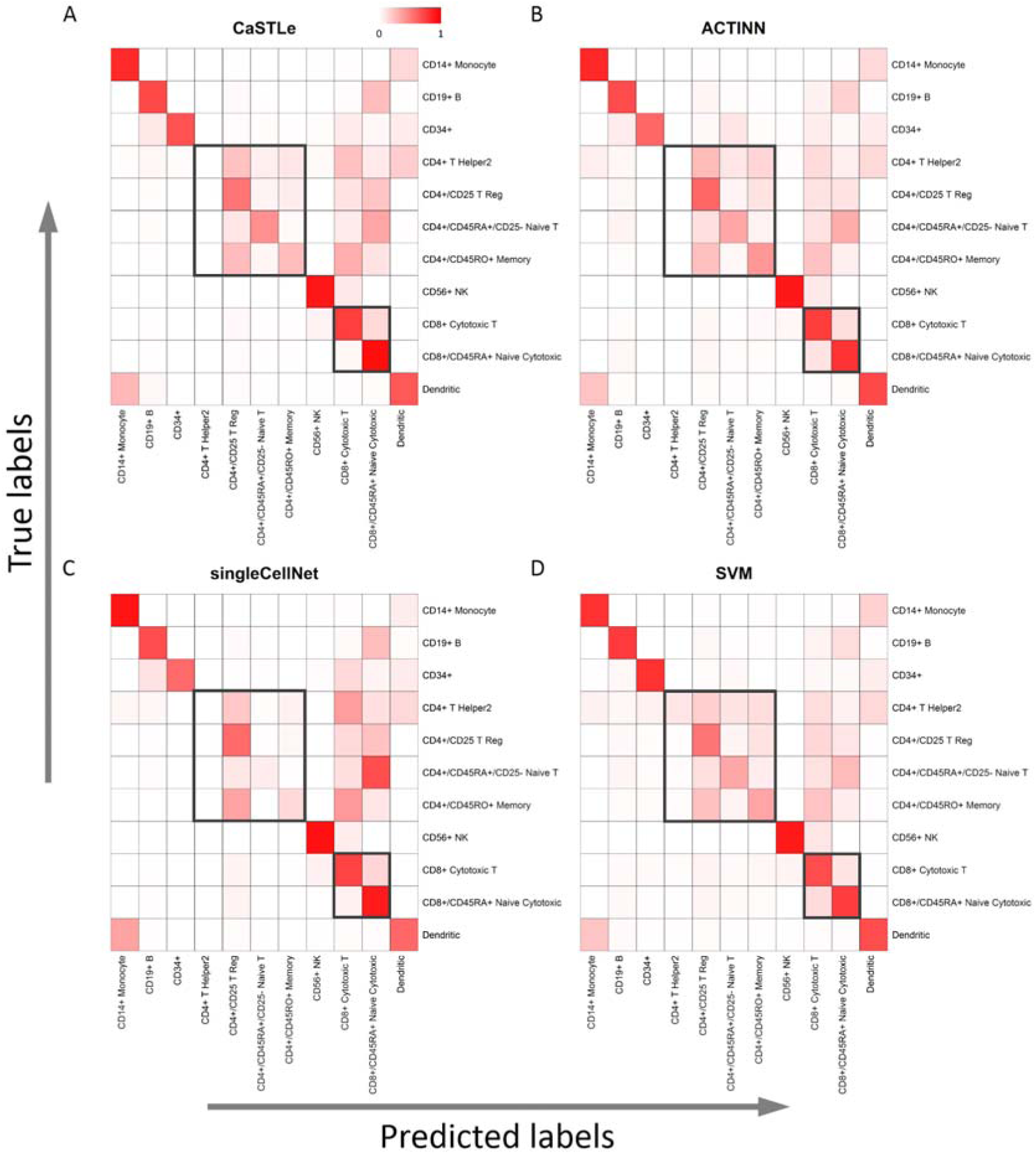
Confusion matrices for the Zheng dataset. Results of the best four classifier **(A)** *CaSTLe*, **(B)** *ACTINN*, **(C)** *singleCellNet*, and **(D)** *SVM* are shown. Rows indicate the true labels and columns indicate the predicted labels. Each cell in the heatmap is colored according to the percentage of overlapping cells between the true and predicted cell population. Black boxes highlight the four subpopulations of CD4 and the two subpopulations of CD8 T-cells.

**Supplementary Figure 2.**
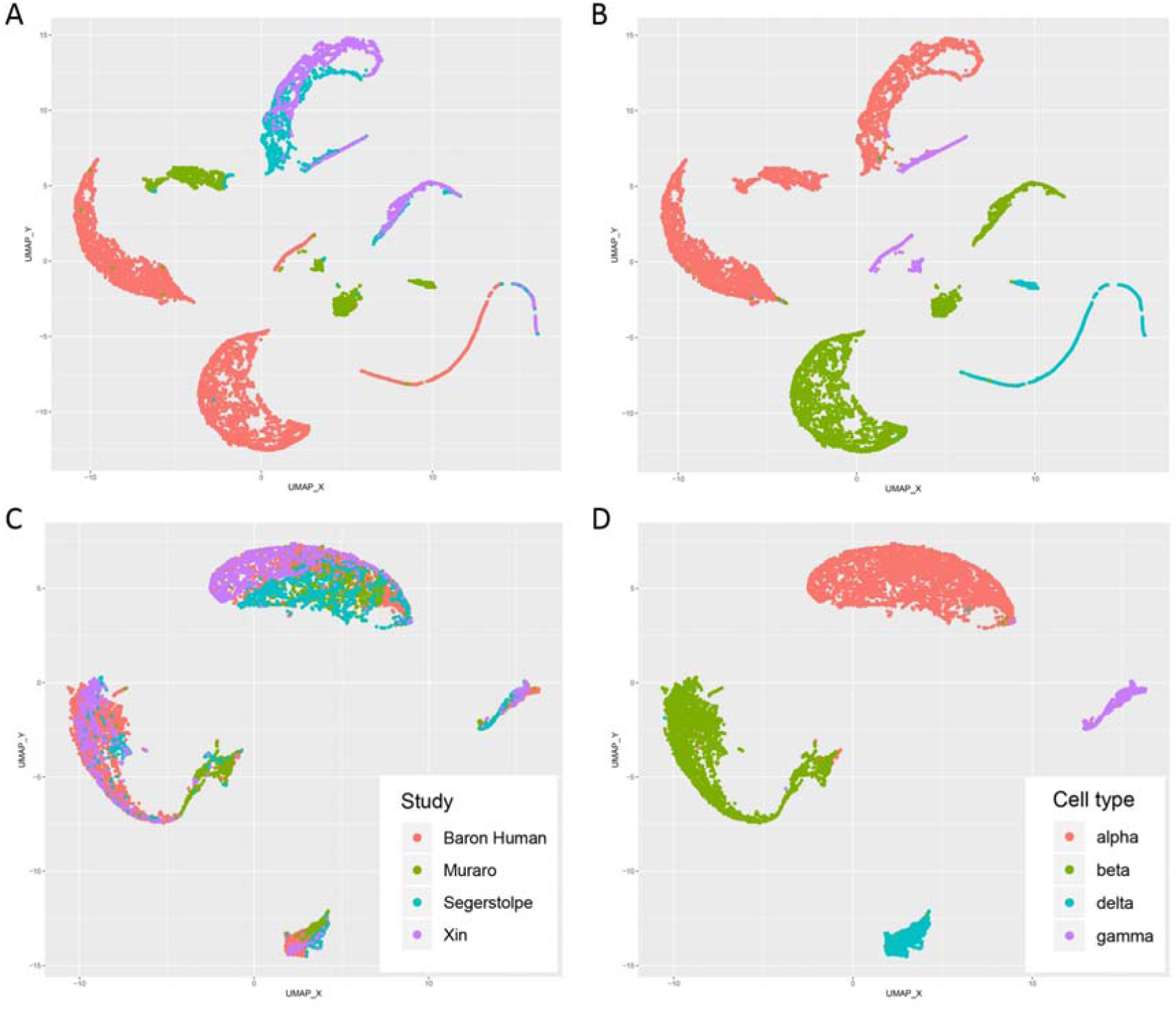
UMAP plots of the four pancreatic datasets used in the across dataset prediction experiment. **(A-B)** UMAP plots before and **(C-D)** after alignment using MNN. In **(A, C)** the cells are colored by dataset and in **(B, D)** the cells are colored by cell type.

